# Adenine-induced kidney disease alters the cortical bone metabolome of C57BL/6J mice in a manner that depends on sex

**DOI:** 10.1101/2025.06.02.657438

**Authors:** Shane Stauffer, Hope Welhaven, Brady Hislop, Bryce A. Jones, Moshe Levi, Ronald K. June, Chelsea M. Heveran

## Abstract

Chronic kidney disease (CKD) increases the likelihood of bone fracture as well as post-fracture mortality. The loss of bone fracture resistance in CKD results from both a loss of bone mass and decreased bone material properties, which together result from changes to the health and activities of bone cells. Determining changes to bone tissue metabolism with CKD may reveal insights important to monitoring and mitigating the decrease in bone fracture resistance that commonly occurs as a result of this disease. In this study, untargeted metabolomics was conducted on marrow-flushed cortical tibiae from female and male C57BL/6J mice fed either a control or 0.2% w/w adenine diet. The diets were continued over 3.5 or 7 weeks to produce different severities of kidney injury. Liquid chromatography mass spectrometry (LC-MS) was used to assess metabolites from tibia extracts. Group comparisons (CKD vs control, 7 weeks vs 3.5 weeks, female vs male) were conducted using principal components analysis (PCA), partial least squares discriminant analysis (PLS-DA), and hierarchical clustering. Clusters of metabolites were also assessed using ensemble clustering and cluster optimization analysis (ECCO).

Volcano plots and VIP scores were used to identify individual metabolites that differed between groups. Pathway analyses were then conducted from these metabolites. The CKD mice, compared with control mice, had dysregulated essential and nonessential amino acid pathways along with altered pathways associated with sugar and fatty acid metabolism. Compared with mice fed an adenine diet for 3.5 weeks, the mice fed an adenine diet over 7 weeks showed dysregulations in the pentose phosphate pathway along with essential and nonessential amino acid metabolism, porphyrin metabolism, steroid hormone biosynthesis, and other pathways relevant to energy production. Sex differences were apparent in the bone tissue metabolomes of females and males. Compared to males, females experienced dysregulations in essential and nonessential amino acid pathways along with other pathways associated with energy derivation, such as pantothenate and CoA biosynthesis. These results demonstrate that CKD alters bone tissue metabolism and reveals novel insights into metabolic dysregulation in disease as well as important sex differences in these metabolic processes.

**Graphical Abstract:** 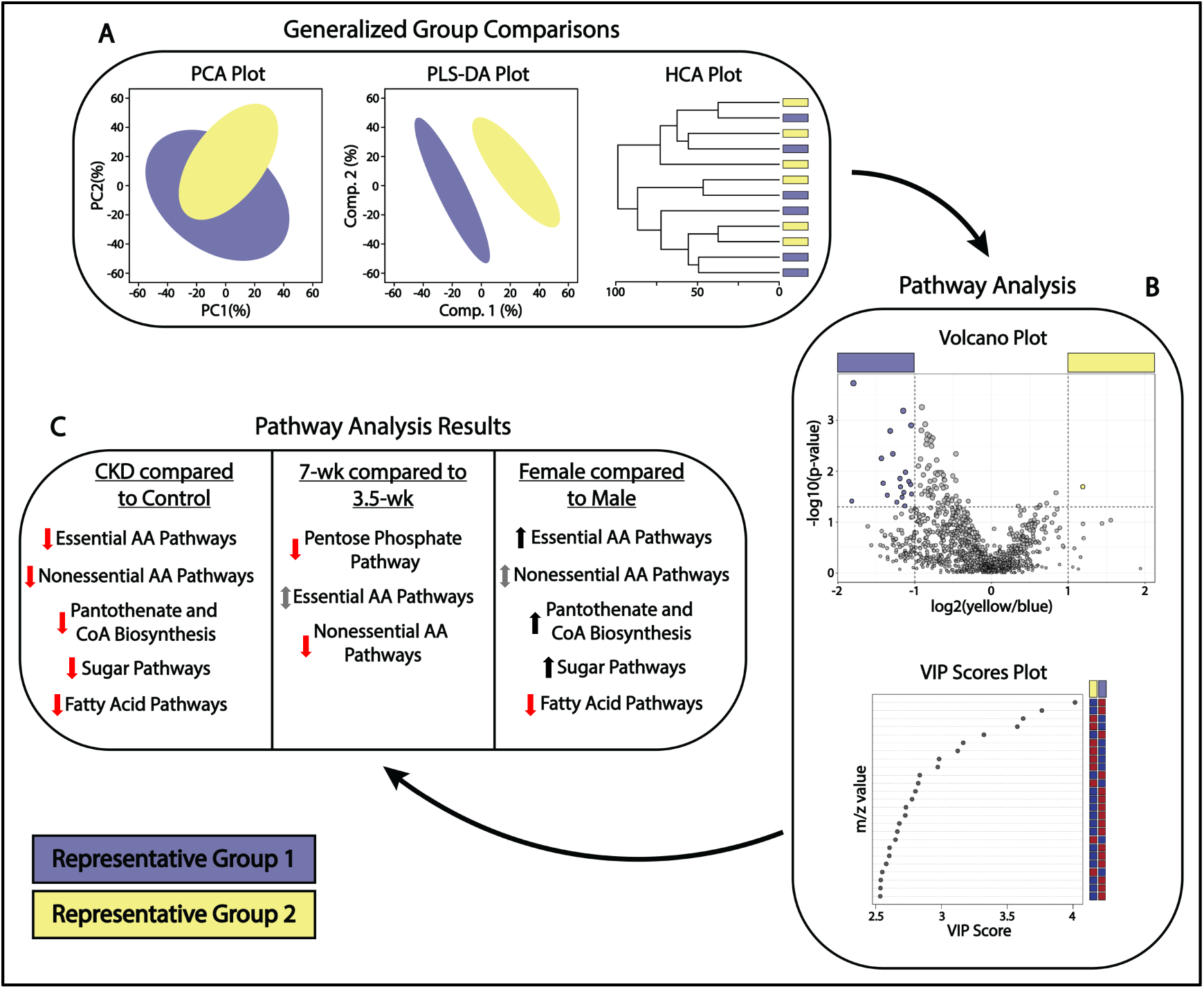

(A) Generalized group comparisons utilizing Partial Component Analysis (PCA), Partial Least Squares-Discriminant Analysis (PLS-DA), and Hierarchical Cluster Analysis (HCA). (B) Pathway analysis utilizing volcano plots and VIP scores plots. (C) Pathway analysis results for comparisons between adenine-induced CKD and control groups, 7-week and 3.5-week diet groups, and female and male groups. A red arrow denotes a set of pathways that were downregulated, a black arrow denotes a set of pathways that were upregulated, and a gray arrow denotes a category of pathways that contained both upregulations and downregulations.

## 1.0 Introduction

Chronic Kidney Disease (CKD) is a major public health issue.^1^ Over 10% (800 million) of the world population and 74% of individuals over the age of 70 suffer from some degree of kidney disease, and its prevalence is expected to increase alongside an aging population.^2–4^ The term Chronic Kidney Disease-Mineral and Bone Disorder (CKD-MBD) has been established to define the abnormalities in mineral/hormone levels (PTH, calcium, phosphate, etc.), bone health, and soft tissue calcification that frequently occur alongside CKD.^4–7^ Within the spectrum of CKD-MBD, renal osteodystrophy (ROD) defines alterations in bone turnover (i.e., formation and resorption).^4,5,8–10^ ROD affects bone mass, microstructure, and matrix properties, which together cause up to a 100-fold increase in fracture prevalence along with a 3-fold increase in post-fracture mortality.^4–6,11^ The detrimental changes to bone tissue and bone cell health in ROD may also be accompanied by altered cellular metabolism.^4–6,8–11^ The metabolic analysis of bone tissue may reveal pathways crucial for understanding bone quality loss associated with CKD and suggest potential biomarkers for ROD detection.

CKD-MBD has the potential to disrupt bone tissue metabolism. Altered metabolism has already been noted in muscle tissue in the context of CKD, with elevated protein synthesis and catabolism, decreased protein anabolism being, and dysregulated glycogenolysis and lipolysis.^12^ It is likely that metabolic dysregulation also occurs in bone tissue. High serum phosphate levels resulting from CKD lead to an increased production of FGF-23 by osteocytes, causing mineral and hormone imbalances that contribute to hyperparathyroidism, altered bone quality, and disrupted bone turnover.^13,14^ CKD also increases oxidative stress, which has been shown to impact bone remodeling and alter bone cell activity.^5,15–17^ Given that 90-95% of cells within the cortical bone are osteocytes, metabolic disruptions to tissue containing these cells can be surveyed through the untargeted metabolomic analysis of osteocyte-enriched (i.e., marrow-removed) bone tissue.^18,19^ This approach has been useful for identifying changes to cortical bone metabolism with altered gut microbiome, aging, and high fat diet.^20–22^

The impact of CKD on bone metabolism has the potential to depend on other factors, such as disease severity. As the stage of CKD progresses, impacts across physiological systems, including the musculoskeletal system, increase.^23–25^ In humans, an increase in CKD stage is also associated with an increase in fracture risk.^4,5,11^ Progressive CKD can be induced in a rodent model via an adenine diet, which leads to the accumulation of urine-insoluble 2,8-dihydroxyadenine in the kidneys, damaging the organs as a result.^26–28^ Prior work demonstrates that an increased duration of the adenine diet leads to an elevation in BUN and plasma creatinine levels, indicating progressive kidney damage.^29^ Therefore, the adenine model is appropriate for comparing the impact of disease severity on bone metabolism.

CKD may also have sex-dependent effects on bone tissue metabolism. In previous studies, mice without CKD showed sex differences in cortical bone metabolism along with difference in bone quality and mechanics. ^21,22,30–34^ Sex differences are also observed in the changes to bone quality associated with the disease. In the adenine model, both female and male adenine-treated mice show decreased cortical bone mineralization and increased cortical porosity, but to different extents, and have divergent effects on cancellous bone (**Table 1**).^28,35–38^ Whether sex difference in bone tissue metabolism differ in the context of adenine-induced CKD remains to be tested.

**Table 1.** The impact of adenine-induced CKD on bone quality measures in C57BL/6 mice.

| Study | Mouse Model | CKD Model | Key Outcomes |
| --- | --- | --- | --- |
| Metzger et. al., 2021, PLOS One | Male and female C57BL/6J mice (n=16 per sex); 26 weeks at euthanasia | 0.2% adenine diet (Male n=8; Female n=8) for 6 weeks from week 16 to week 22 | **Male Aden. vs Ctrl (femurs):**<br>• 27% ↓ BV/TV %<br>• 31% ↓ Cortical Bone Thickness<br>• 22% ↓ Cortical Bone Area<br>• High Bone Turnover<br>**Female Aden. vs Ctrl (femurs):**<br>• 97% ↑ BV/TV %<br>• 21% ↓ Cortical Bone Thickness<br>• Minimal change in Cortical Bone Area<br>• High Bone Turnover |
| Metzger et. al., 2021, Bone | Female C57BL/6J mice (n=16); 26 weeks at euthanasia | 0.2% adenine diet (n=8) for 6 weeks from week 16 to week 22 | **Aden. vs Ctrl (femurs):**<br>• 97% ↑ BV/TV %<br>• 21% ↓ Cortical Bone Thickness<br>• No difference in cortical bone area<br>• 2900% ↑ in Cortical Porosity<br>• High Bone Turnover |
| Metzger et. al., 2024, JBMR Plus | Male C57BL/6J mice (n=48); 28 weeks at euthanasia | 0.2% adenine diet (n=12) for 6 weeks from week 16 to week 22 | **Aden. vs Ctrl (femurs):**<br>• 45% ↓ BV/TV %<br>• 26% ↓ Cortical Bone Thickness<br>• 19% ↓ Cortical Bone Area<br>• 434% ↑ Cortical Porosity<br>• 19% ↓ Trabecular Thickness<br>• 28% ↑ Trabecular Separation<br>• High bone turnover |
| Tani et. al., 2017, Scientific Reports | Male C57BL/6J mice (n=35); 14 (n=10), 16 (n=5), 18 (n=5), or 20 (n=15) weeks at euthanasia | 0.2% adenine diet for 6 (n=5) or 12 (n=5) weeks from week 8 to week 14 or 20, respectively | **6-week Aden. diet vs Ctrl (femurs):**<br>• Evident kidney atrophy with regions of excess collagen growth<br>• No change in BV/TV<br>• 19% ↓ BMD<br>• 20% ↓ Cortical Mineral Density<br>• 27% ↓ Cortical Mineral Volume<br>**12-week Aden. diet vs Ctrl (femurs):**<br>• Increased levels of kidney atrophy and regions of excess collagen growth (compared to 6-week diet)<br>• No change in BV/TV<br>• 11% ↓ BMD<br>• 19% ↓ Cortical Mineral Density |
|  |  |  | • 30% ↓ Cortical Mineral Volume |
| Damrath et. al., 2022, Bone | Female C57BL/6J mice (n=49); 24 weeks at euthanasia<br>Femurs, tibiae, and L4 vertebrae examined | 0.2% adenine diet for 6 weeks from week 14 to week 20 | **Aden. vs Ctrl (tibiae):**<br>• 14% ↑ BV/TV %<br>• 25% ↓ Cortical Bone Thickness<br>• 8% ↓ Trabecular Thickness<br>• 8% ↓ Trabecular Separation<br>• 20% ↓ Bone Area/Total Area %<br>• 11% ↑ in Tibial Yield Stress<br>• 22% ↓ in Tibial Ultimate Stress<br>• No change in bone turnover |

The purpose of this study was to test the hypothesis that adenine-induced CKD dysregulates the cortical bone metabolome, and that the specific disruptions depend on CKD severity and sex. This hypothesis was tested through untargeted assessment of marrow-flushed tibiae from male and female C57BL/6 mice fed control or adenine diets over two lengths of time (3.5 weeks and 7 weeks).

## 2.0 Materials and Methods

### 2.1 Animal Model

C57BL/6J male/female mice were purchased from The Jackson Laboratory (Bar Harbor, ME). Mice were fed a standard chow (D19120401i, Research Diets, New Brunswick, NJ) until 12 weeks of age.

Adenine-treated mice (males and females) were then given a diet enriched with 0.2% w/w adenine for either 3.5 or 7 weeks (D19120402i, Research Diets), with the respective controls being given a standard diet over those periods. The mice were euthanized via carbon dioxide. At the study endpoint, the experimental groups included 15.5-week males (Ctrl: n=6; CKD: n=6), 15.5-week females (Ctrl: n=6; CKD: n=6), 19-week males (Ctrl: n=6; CKD: n=4), and 19-week females (Ctrl: n=6; CKD: n=6) (**Table 2**). All mice were administered a saline vehicle as a control for another study. Animal procedures were approved by the Georgetown University IACUC.

**Table 2.**
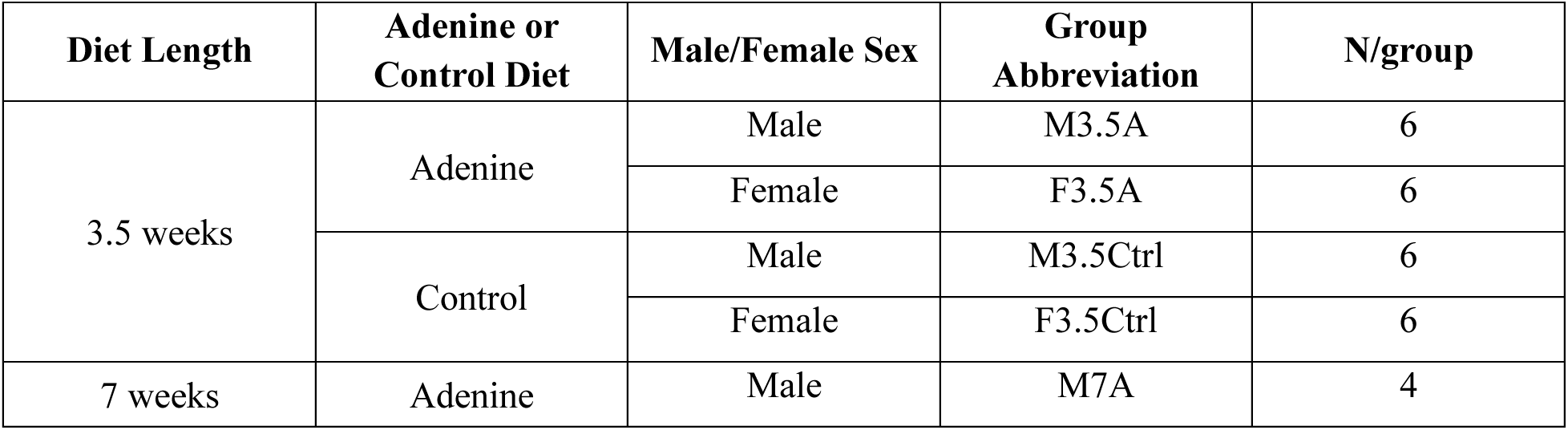
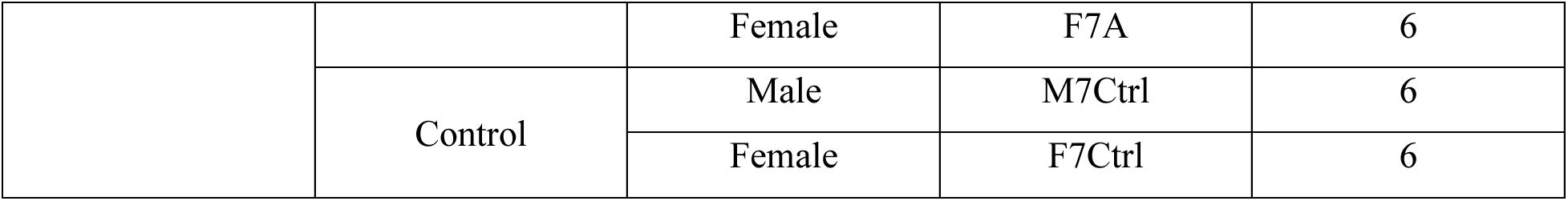
Study design.

Plasma was separated from heparinized blood collected at euthanasia and was assessed for blood urea nitrogen (BUN) and creatinine levels, both markers of kidney disease severity. Plasma creatinine was quantified with a colorimetric kit (DICT-500, BioAssay Systems), and BUN was quantified as previously described.^39^ The values from the M7Ctrl and M7A groups have been previously reported.^40^

### 2.2 Preparation of Cortical Bone for LC-MS

The hind limbs were frozen prior to biochemical analysis, and 8 groups (total n = 46) with differing diets, diet durations, and sexes were utilized to compare the metabolomes of isolated cortical bone (**Table 2**). Mouse tibiae were isolated via a standard dissection protocol. The distal and proximal ends of the tibiae were then removed, and the marrow within the cortical shaft was removed via nested centrifugation. A razor blade was used to remove the tip of a 0.5-mL microcentrifuge tube so that the hole was smaller than the width of the tibiae. The microcentrifuge tube was placed in a 1.5-mL microcentrifuge tube, and a tibiae and 0.25 mL of PBS was added to the smaller centrifuge tube. The unit was then centrifuged at 15,000 x g for 1 minute, leaving a flushed cortical shaft. Cortical shafts were isolated for all mouse samples (n = 46) in the study. The cortical bone samples were wrapped in aluminum foil and were stored at-80°C until metabolites were extracted.

Bone metabolites were extracted from isolated cortical bone by freezing the samples in liquid nitrogen for 2 hours before crushing the samples with a hammer to maximize bone surface area. The crushed bone was suspended in 70:30 methanol:acetone and metabolites were extracted via five-cycles of 1 minute of vortexing followed by 4 minutes of macromolecule precipitation in a-20°C freezer. The samples were then stored in a-20°C freezer overnight to maximize precipitation. Samples were then centrifuged the next day at 16,000 x g for 10 minutes (4°C) to separate the metabolite suspension from the remaining cortical bone/other material. The supernatants were extracted and dried via vacuum concentration. Metabolites were resuspended in 50:50 water:acetonitrile and transferred to MS snap cap tubes prior to LC-MS runs (**Figure 1**). All solvents used were HPLC grade or higher.

**Figure 1.**
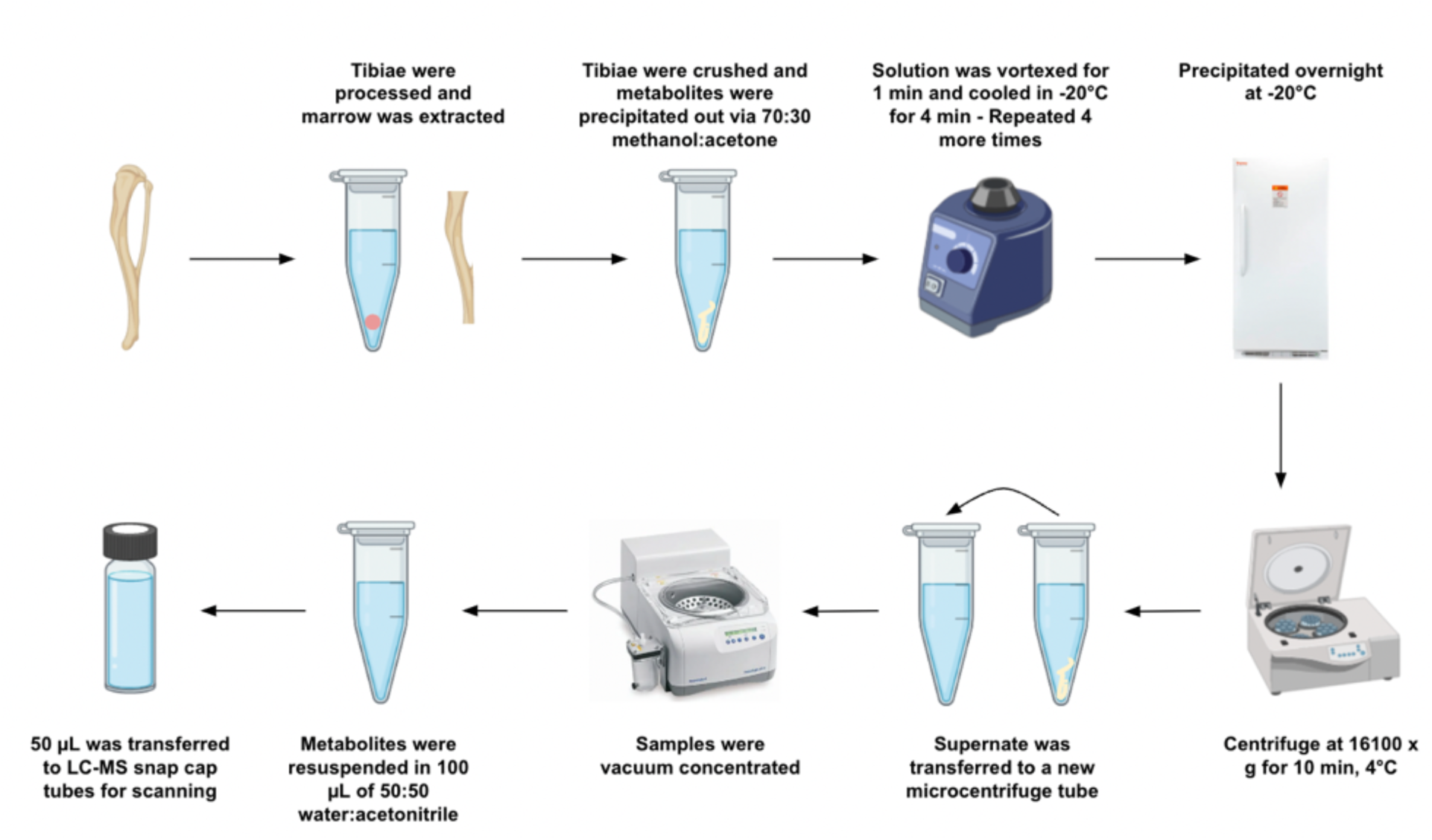
Process for extracting metabolites from the cortical bone of mouse tibiae.

**Figure 2.**
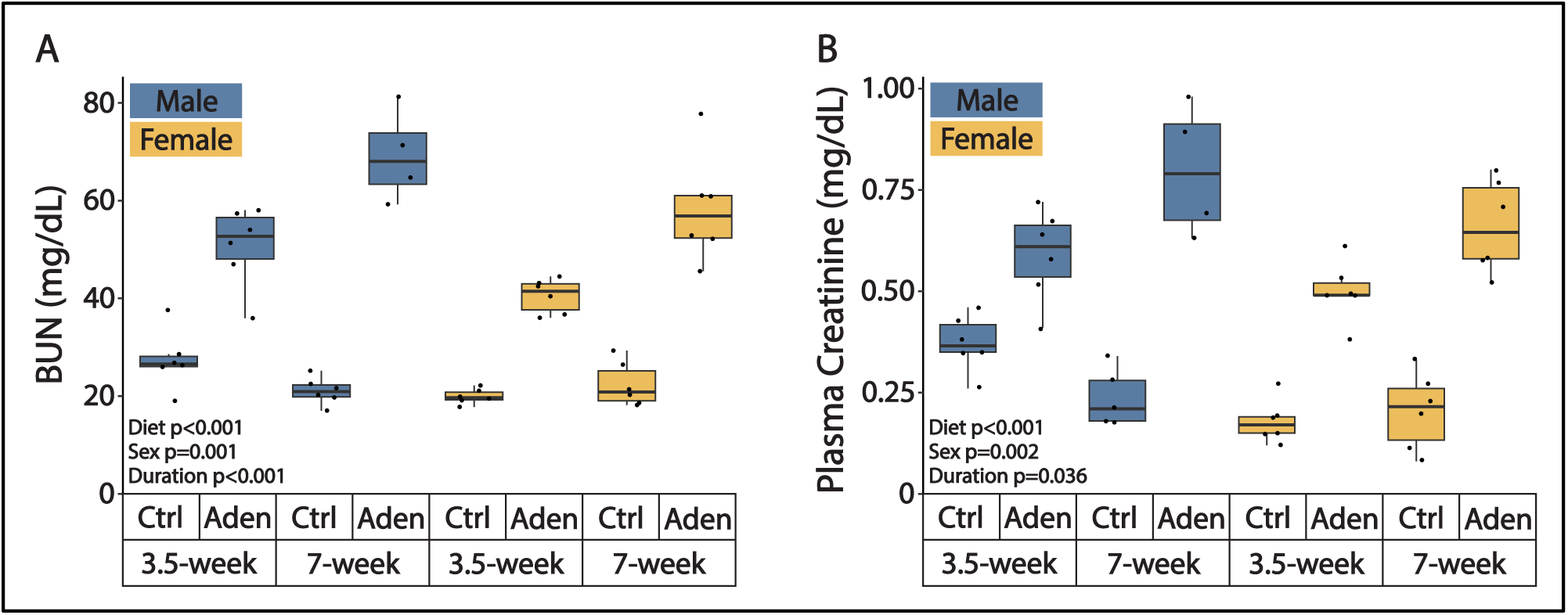
The adenine diet induces kidney disease in a time-dependent manner in both sexes. (A) BUN levels for male and female mice fed control or adenine diets for 3.5 and 7 weeks. (B) Plasma creatinine levels for male and female mice fed control or adenine diets for 3.5 and 7 weeks. Data are means ± SEM, n = 4-6 mice per group.

### 2.3 Mass Spectrometry

All isolated cortical bone samples were then analyzed via liquid chromatography-mass spectrometry (LC-MS) using an Agilent 1290 LC coupled to an Agilent 6538 QTOF mass spectrometer in positive mode (Agilent Technologies, Santa Clara, CA, USA). A Cogent Diamond Hydride HILIC column (2.2 μM, 120 Å, 150 mm x 2.1 mm; MicroSolv, Leland, NC, USA) was used for chromatography. 5 μL of each sample was injected (including blank controls), and blank samples were analyzed every 10 experimental samples for quality control. Peak intensity values were identified and exported for m/z values using Agilent Masshunter Qualitative Analysis software. XCMS was then used to export LC-MS data and convert it for analysis. Statistical analysis of mass features was completed using MetaboAnalyst, with the Kyoto Encyclopedia of Genes and Genomes (KEGG) being utilized to confirm retention time and the exact mass of detectable metabolite features.

### 2.4 Statistical Analysis

#### ANOVA analysis for plasma biomarkers

Blood urea nitrogen and plasma creatinine were compared across groups using 3-way ANOVA, with the factors of diet (adenine versus control), duration (3.5 vs 7 weeks), or sex (male vs female) and their interactions. Residuals were checked for normality and equal variance. Analyses were performed in Minitab v19.

#### Group comparisons

Metabolomics data was analyzed with Metaboanalyst (https://www.metaboanalyst.ca/). Using established procedures, raw data was log transformed, standardized, and auto-scaled (mean-centered divided by standard deviation per variable).^41^ 40% of features were filtered out based on standard deviation to minimize background noise and remove inconsistently detected metabolites. Group comparisons included t-tests, fold change analysis, hierarchical cluster analysis (HCA), principal component analysis (PCA), partial least-squares discriminant analysis (PLS-DA), and volcano plot analysis. HCA and PCA are methods of unsupervised analysis that examine differences between experimental groups, with HCA analyses identifying sub-groups based on metabolic profiles. PLS-DA is a supervised method that examines data variation based on the sample groupings provided by the user.

#### Ensemble clustering combined with cluster optimization

Metabolomic profiles were analyzed using ensemble clustering combined with cluster optimization (ECCO) to identify metabolites which consistently co-cluster. Ensemble clustering combines the solutions from multiple clustering algorithms together to determine the metabolites which responds in a similar manner to an experimental group (here, CKD, diet duration (i.e., severity), or sex).^42^ A total of 4 linkage functions and 10 distance functions were used, with this “ensemble” allowing assessment of which metabolic features are most often grouped together. To identify the optimal number of clusters, clustering optimization was performed prior to the use of ensemble clustering. By combining the “optimal” clustering solutions with ensemble clustering, this approach increased the likelihood of detecting the natural groupings in the data.

#### Pathway identification

Volcano and variable importance in projection (VIP) plot analyses were used to explore differentially regulated metabolite features within the study. Volcano plots compare metabolic features expressed within a specific comparison and determine significant features based on both fold change and p-value. Fold change compares the median intensity of a specific metabolite in one group and compares it to the intensity of the metabolite in the other group, judging significance based on magnitude of difference.

Volcano plots were analyzed using a fold change value of 2 and a p-value of 0.05. False discovery rate (FDR) correction was applied to the p-values associated with the volcano plots to account for multiple comparisons. VIP plots are created from the PLS-DA data and describe metabolite features that contribute the most to differences within compared groups. The metabolites found in both comparisons are then subject to pathway analysis. Pathways are determined using MetaboAnalyst’s MS Peaks to Pathways feature with the mummichog algorithm. Metabolites are compared to the *Mus musculus* [KEGG] pathway library and features are matched to pathways within this library.

## 3.0 Results

### 3.1 Plasma biomarkers confirm kidney disease resulting from adenine diet

A 3-way ANOVA was performed to test whether sex, diet type, and diet duration or their interactions affect BUN and creatinine. BUN was significantly increased with adenine diet (+ 135%, p = <0.001), male sex (+12%, p = 0.001), and diet duration (+17%, p = <0.001) (**Supplementary Information – Table S.1**). One BUN outlier was excluded due to technical error. Creatinine was also increased with adenine diet (+121%, p = <0.001), male sex (+32%, p = 0.002), and diet duration (+18%, p = 0.036) (**Supplementary Information – Table S.2**). There were no significant two-or three-way interactions between diet, sex, and diet duration for creatinine or BUN.

### 3.2 Group-level differences in cortical bone metabolism: adenine-induced CKD vs control

The strategy for metabolic analyses was to 1) assess whether clusters of metabolites were dysregulated between groups (ECCO analysis), 2) assess group-level differences in metabolism using HCA, PCA, and PLS-DA analyses, 3) assess whether individual metabolites are dysregulated using volcano and VIP analyses, and 4) conduct pathway analyses from dysregulated metabolites. This strategy was employed for analyses of the effects of CKD (vs control), CKD severity (3.5 vs 7 weeks of adenine diet), and male vs female sex.

First, ECCO was used to perform clustering analyses for all samples (i.e., metabolites of all mice) in the study. The clustering solutions derived from ECCO were not strongly aligned with factors in this study (i.e., CKD, severity, sex) (**Supplementary Information – Figure S.1**). Next, differences in metabolism between adenine and control groups were analyzed when pooled across sex and severity groups (all adenine mice vs all control mice). 40.8% and 26.4% of variation was accounted for in the first 2 dimensions of PCA and PLS-DA, respectively (**Figure 3A, B**). The volcano plot for the pooled adenine vs control comparison did not reveal significant metabolic differences between these groups (**Figure 3C**).

**Figure 3.**
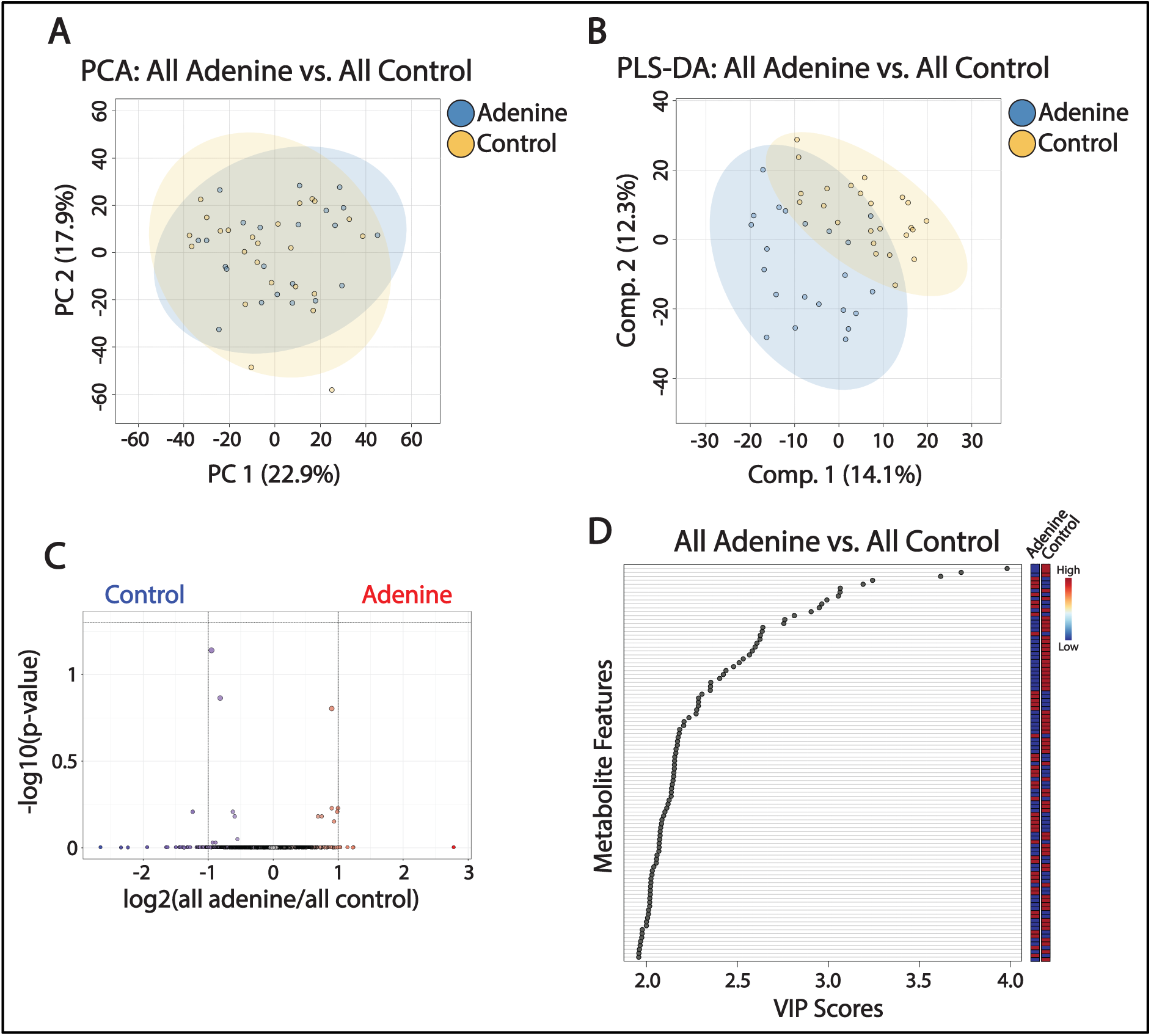
Outputs from aggregated comparisons of CKD presentation. (A) PCA plot comparing all adenine and control samples. (B) PLS-DA plot comparing all adenine and control samples. (C) Volcano plot comparing all adenine control samples. (D) Top 100 VIP scores plot for the comparison between all adenine and control samples.

VIP scores were then used to define the metabolites that contributed the most to the differences between compared groups based on the 100 highest VIP scores representing the metabolites with the greatest dysregulation between the groups (adenine vs control). A VIP score of 1 or greater indicates that the metabolic feature is significantly important to the differences between the compared groups.^43^ For the pooled adenine vs control mice comparison, the top 100 metabolites were found to have VIP scores between 1.96 and 3.98 (**Figure 3D**). From pathway analysis, the metabolites were consistent with downregulated fatty acid and sugar pathways (galactose, starch, and sucrose metabolism), other pathways associated with energy metabolism (porphyrin metabolism, ubiquinone and other terpenoid-quinone metabolism, pantothenate and CoA biosynthesis), and essential (valine, leucine, and isoleucine biosynthesis and degradation, lysine degradation, and methionine, threonine, and histidine metabolism) and nonessential (arginine, proline, glycine, serine, cysteine, and tyrosine metabolism) amino acid pathways in mice fed an adenine diet (**Table 3**).

**Table 3.** Metabolic pathways identified by the comparison between mice with and without adenine-induced CKD (pooled males and females). VIP scores labeled with an asterisk indicate the pathways that were most significantly dysregulated in their associated comparisons.

| Pathway | Comparison | Pathway Total | Pathway Hits | VIP Score | Trends |
| --- | --- | --- | --- | --- | --- |
| Fatty Acid Degradation | All Aden vs Ctrl | 39 | 5 | 2.16 | ↓ in Aden |
| Fatty Acid Elongation | All Aden vs Ctrl | 39 | 3 | 2.16 | ↓ in Aden |
| Porphyrin Metabolism | All Aden vs Ctrl | 31 | 5 | 2.6 | ↓ in Aden |
| Steroid Hormone Biosynthesis | All Aden vs Ctrl | 79 | 48 | 2.17 | ↓ in Aden |
| Purine Metabolism | All Aden vs Ctrl | 71 | 4 | 2.63 | ↓ in Aden |
| Galactose Metabolism | All Aden vs Ctrl | 27 | 5 | 2.14 | ↓ in Aden |
| Starch and Sucrose Metabolism | All Aden vs Ctrl | 15 | 4 | 2.14 | ↓ in Aden |
| Pantothenate and CoA Biosynthesis | All Aden vs Ctrl | 20 | 3 | 3.98* | ↓ in Aden |
| Tyrosine Metabolism | All Aden vs Ctrl | 42 | 17 | 2.78 | ↓ in Aden |
| Arginine and Proline Metabolism | All Aden vs Ctrl | 36 | 8 | 3.98* | ↓ in Aden |
| Glycine, Serine, and Threonine Metabolism | All Aden vs Ctrl | 34 | 3 | 3.98* | ↓ in Aden |
| Lysine Degradation | All Aden vs Ctrl | 30 | 6 | 3.98* | ↓ in Aden |
| Histidine Metabolism | All Aden vs Ctrl | 16 | 4 | 2.51 | ↓ in Aden |
| Cysteine and Methionine Metabolism | All Aden vs Ctrl | 33 | 7 | 2.63 | ↓ in Aden |
| Valine, Leucine, and Isoleucine Biosynthesis | All Aden vs Ctrl | 8 | 6 | 3.98* | ↓ in Aden |
| Valine, Leucine, and Isoleucine Degradation | All Aden vs Ctrl | 40 | 9 | 3.98* | ↓ in Aden |

### 3.3 The effect of adenine-induced CKD on cortical bone metabolism depends on sex

The impacts of adenine-induced CKD on cortical bone metabolism were compared for males and females. Comparisons were first performed between adenine (females and males) and control (females and males) groups, with PCA and PLS-DA accounting for 40.8% and 20.6% of variation, respectively (**Figure 4A,B**). Neither comparison detected significantly dysregulated features when analyzed using volcano plots (**Figure 4E, F**).

**Figure 4.**
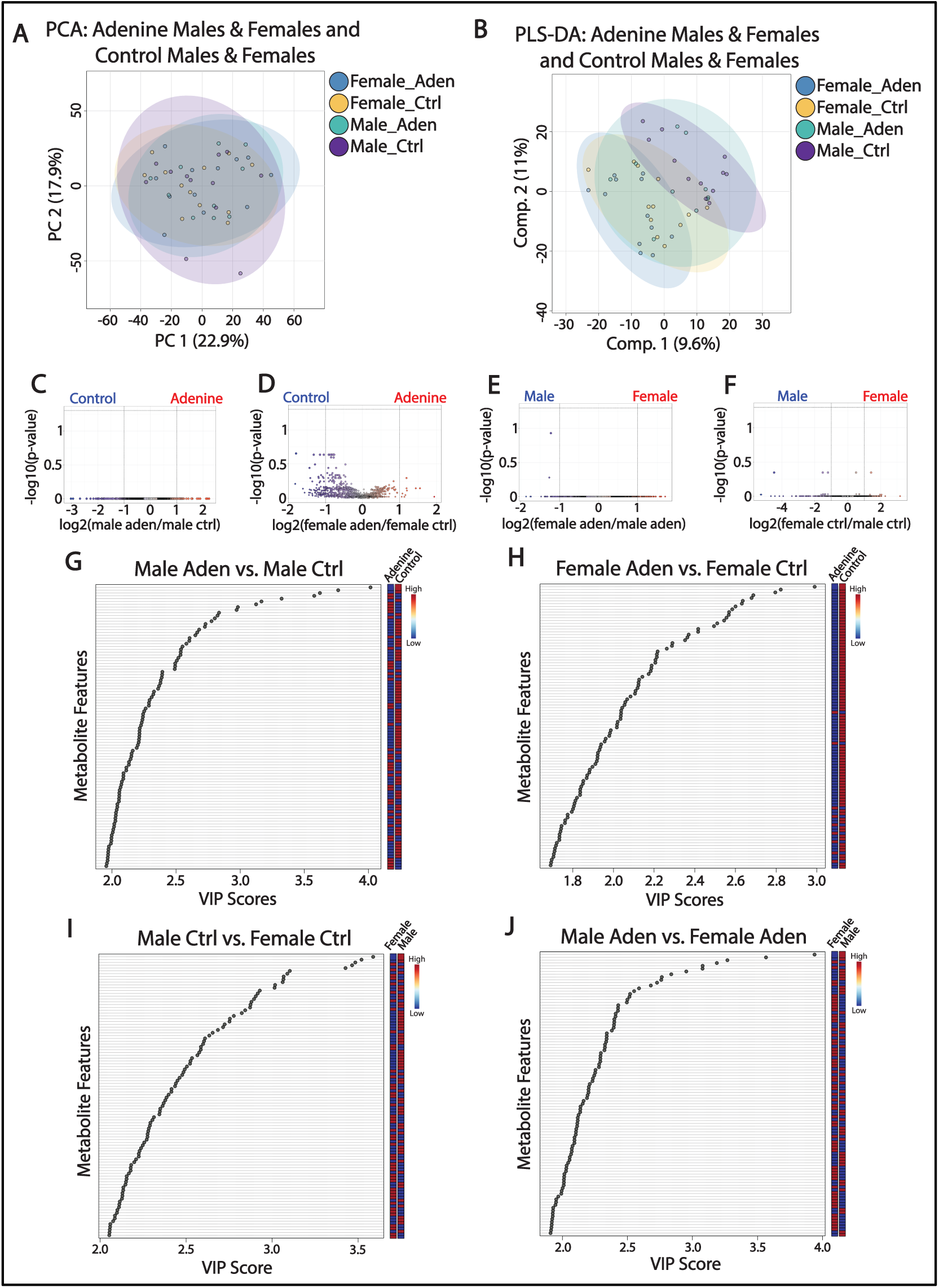
Outputs from the expanded comparisons of male/female sex. (A) PCA plot comparing adenine females, adenine males, control females, and control males. (B) PLS-DA plot comparing adenine females, adenine males, control females, and control males. (C) Volcano plot comparing adenine males to control males. (D) Volcano plot comparing adenine females to control females. (E) Volcano plot comparing adenine females to adenine males. (F) Volcano plot comparing control females to control males. (G) Top 100 VIP scores plot for the comparison between adenine and control males. (H) Top 100 VIP scores plot for the comparison between adenine and control females. (I) Top 100 VIP scores plot for the comparison between control males and females. (J) Top 100 VIP scores plot for the comparison between adenine males and females.

For male versus female control mice, the VIP scores for the top 100 metabolites ranged from 2.05 to 3.59 (**Figure 4I**). Female mice experienced downregulated fatty acid degradation, steroid hormone biosynthesis, tyrosine metabolism, and ubiquinone and other terpenoid-quinone biosynthesis compared to males (**Table 4**). Female mice also experienced upregulated pantothenate and CoA biosynthesis, galactose metabolism, and essential (lysine degradation, threonine metabolism, and valine, leucine, and isoleucine biosynthesis and degradation) and nonessential (arginine, proline, glycine, and serine metabolism) amino acid pathways compared to male mice (**Table 4**).

**Table 4.**
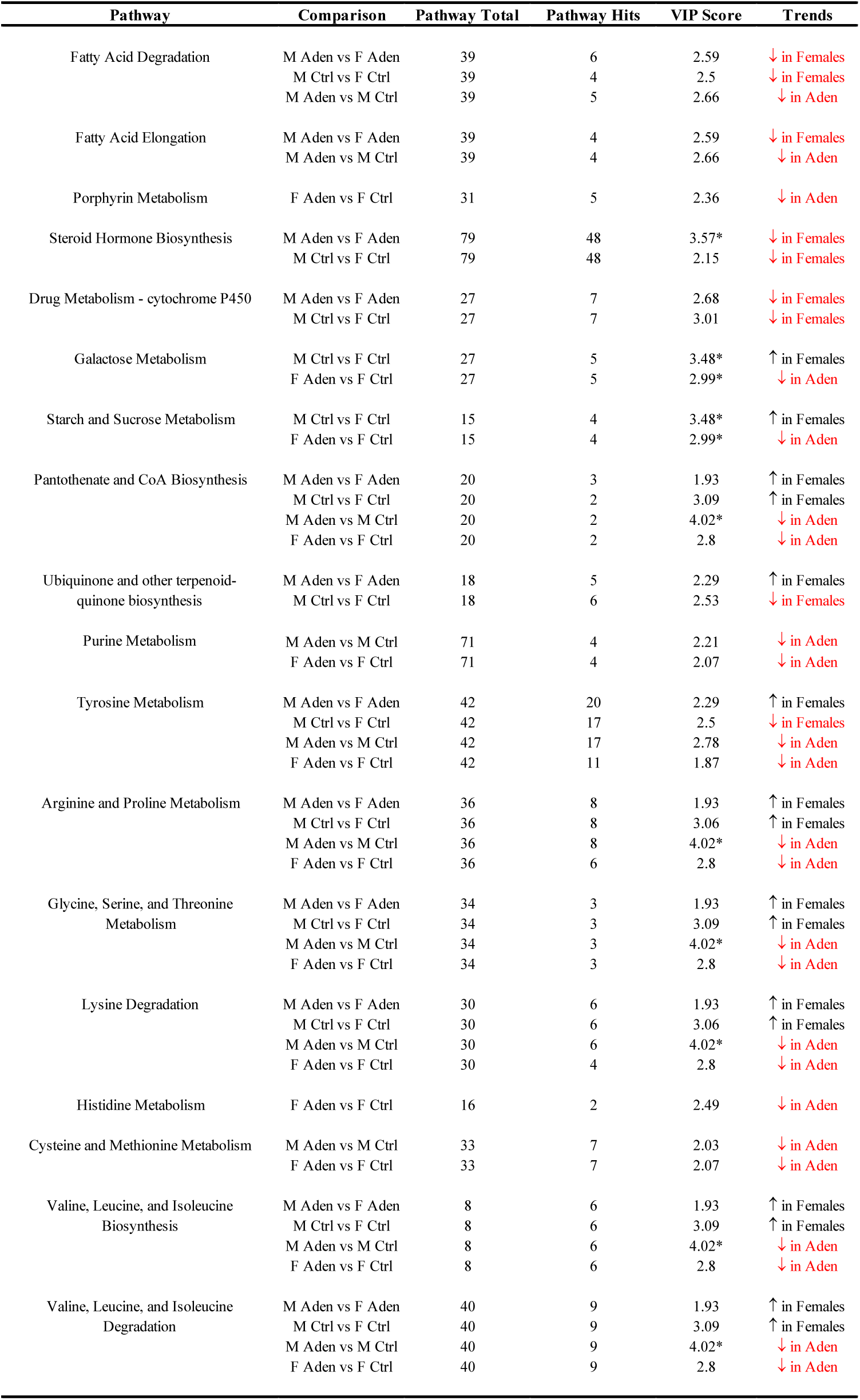
Metabolic pathways identified for comparisons between male and female mice with and without adenine-induced CKD. VIP scores labeled with an asterisk indicate the pathways that were most significantly dysregulated in specific comparisons.

For male versus female adenine mice, the VIP scores for the top 100 metabolites ranged from 1.91 to 3.94 (**Figure 4J**). Female mice experienced downregulated fatty acid degradation and elongation, steroid hormone biosynthesis, and drug metabolism – cytochrome P450 compared to males (**Table 4**). Females also experienced upregulated pantothenate and CoA biosynthesis, ubiquinone and other terpenoid-quinone biosynthesis, and essential (lysine degradation, threonine metabolism, and valine, leucine, and isoleucine biosynthesis and degradation) and nonessential (arginine, proline, glycine, serine, and tyrosine metabolism) amino acid pathways compared to male mice (**Table 4**).

The metabolite dysregulation experienced in the male/female sex comparisons were very similar between adenine and control comparisons. However, tyrosine metabolism was downregulated in female control mice but was upregulated in female adenine mice. Moreover, ubiquinone and other terpenoid-quinone biosynthesis was downregulated in female control mice while being upregulated in female adenine mice.

Metabolomic profiles were then compared between female adenine and control mice, and between male adenine and control mice. PCA and PLS-DA accounted for 40.8% and 20.6% of variation, respectively, in each comparison (**Figure 4A, B**). No significant metabolites were identified from the volcano plots of either comparison (**Figure 4C, D**). Significant metabolites were identified from top 100 VIP scores plots for both comparisons, though (**Figure 4G, H**). The range of VIP scores for the top 100 metabolites in the female adenine and control mice comparison was 1.96-4.02. For the comparison between male adenine and control mice, the range of VIP scores was 1.69-2.99.

Pathway analysis showed that female adenine mice experienced downregulated pantothenate and CoA biosynthesis, essential (valine, leucine, and isoleucine biosynthesis and degradation, lysine degradation, methionine metabolism, and threonine metabolism) and nonessential (arginine and proline metabolism, glycine and serine metabolism, cysteine metabolism, and tyrosine metabolism) amino acid pathways, and purine metabolism compared to their respective controls; very similar pathway alterations were seen for male adenine versus control mice (**Table 4**). The comparison between female adenine and control mice, and the comparison between male adenine and control mice, differed in that fatty acid pathways were only dysregulated in male adenine mice (downregulated compared to control diet mice), while sugar pathways, porphyrin metabolism, and histidine metabolism were only dysregulated in female adenine mice (downregulated compared to control diet mice) (**Table 4**).

### 3.4 The effect of adenine-induced CKD on cortical bone metabolism depends on disease severity

The impacts of diet length on bone metabolism alterations were then assessed. Comparisons between adenine (3.5 weeks vs 7 weeks of adenine diet) and control (3.5 weeks vs 7 weeks of control diet) diet mice were conducted. The latter comparison provides an estimate of baseline metabolism differences over the diet duration (e.g. reflects aging and potentially other factors not attributable to kidney disease). 40.8% and 32.7% of variation was accounted for in the PCA and PLS-DA plots for these comparisons, respectively (**Figure 5A, B**).

**Figure 5.**
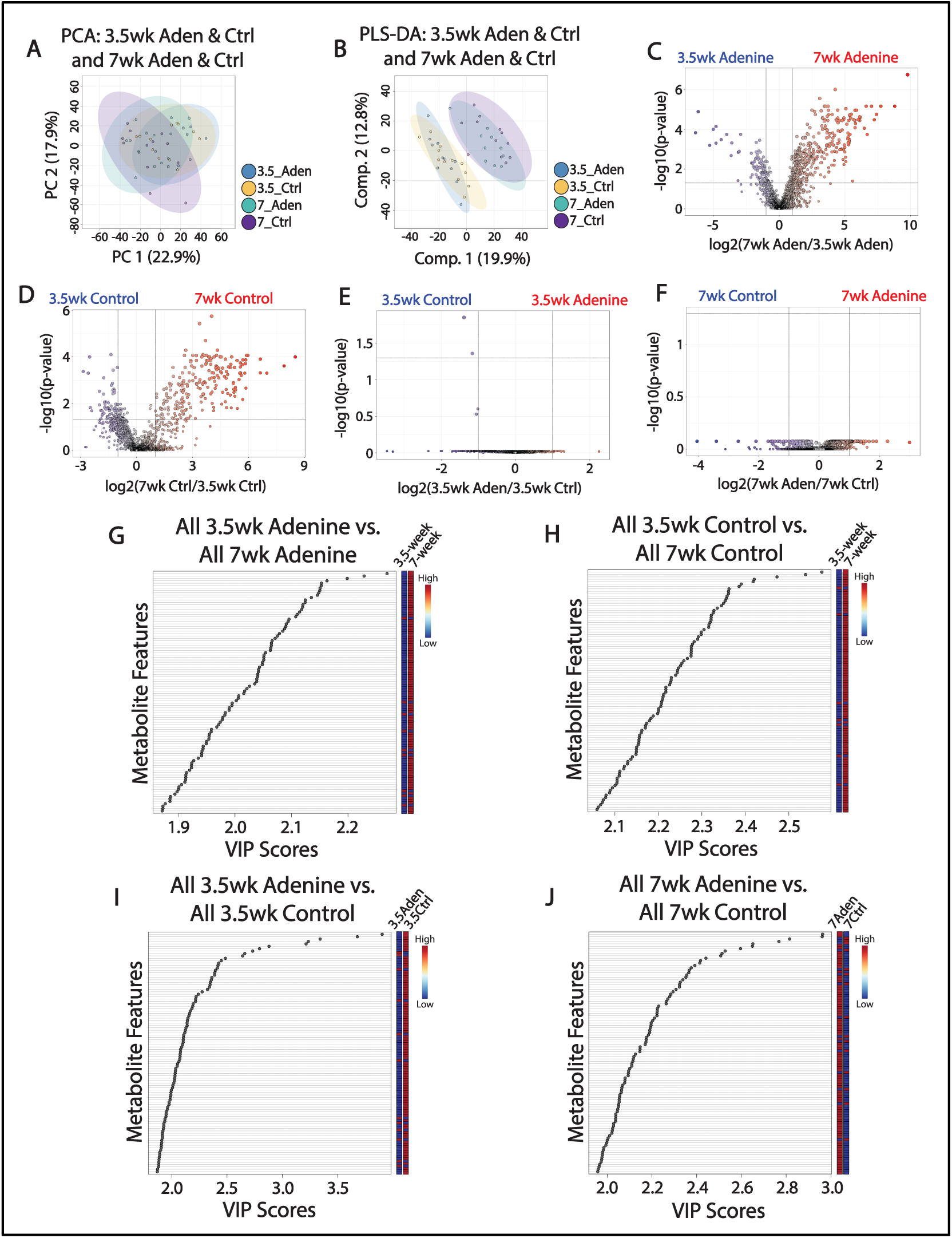
Outputs from the expanded comparisons of male/female sex. (A) PCA plot comparing mice fed a 3.5-week control diet, a 3.5-week adenine diet, a 7-week control diet, and a 7-week adenine diet. (B) PLS-DA plot comparing mice fed a 3.5-week control diet, a 3.5-week adenine diet, a 7-week control diet, and a 7-week adenine diet. (C) Volcano plot comparing mice fed a 7-week adenine diet to mice fed a 3.5-week adenine diet. (D) Volcano plot comparing mice fed a 7-week control diet to mice fed a 3.5-week control diet. (E) Volcano plot comparing mice fed a 3.5-week adenine diet to those fed a 3.5-week control diet. (F) Volcano plot comparing mice fed a 7-week adenine diet to those fed a 7-week control diet. (G) Top 100 VIP scores plot for the comparison between mice fed an adenine diet over 3.5 or 7 weeks. (H) Top 100 VIP scores plot for the comparison between mice fed a control diet over 3.5 or 7 weeks. (I) Top 100 VIP scores plot for the comparison between mice fed a 3.5-week adenine diet or 3.5-week control diet. (J) Top 100 VIP scores plot for the comparison between mice fed a 7-week adenine diet or 7-week control diet.

Volcano plot analysis detected significant metabolic features. In comparing pooled groups of mice fed either a 3.5-or 7-week adenine diet, 495 metabolites were determined to be significant. Of these, 409 were upregulated in 7-week adenine mice and the remaining 86 were downregulated (**Figure 5C**). From pathway analysis, it was found that 7-week adenine mice experienced downregulated pentose phosphate pathway, fatty acid degradation, and some essential (methionine metabolism) and nonessential (cysteine metabolism) amino acid pathways (**Table 5B**). Upregulations in pathways associated with energy metabolism (porphyrin metabolism, ubiquinone and other terpenoid-quinone biosynthesis) were also observed in 7-week adenine mice (**Table 5B**).

**Table 5.** Metabolic pathways identified through comparisons of 3.5 or 7 weeks of adenine or control diets. (A) Pathways identified from top 100 VIP scores plots for each of the relevant comparisons. (B) Pathways identified from Volcano plots for each of the relevant comparisons. VIP scores with an asterisk indicate the pathways that were most significantly dysregulated in specific comparisons.

| Pathway | Comparison | Pathway Total | Pathway Hits | VIP Score | Trends |
| --- | --- | --- | --- | --- | --- |
| **(A) Pathways of interest from top 100 VIP scores** |  |  |  |  |  |
| Amino sugar and nucleotide sugar metabolism | M3.5Ctrl_M7Ctrl | 42 | 2 | 2.22 | ↑ in 7wk |
| Pentose Phosphate Pathway | 3.5A_7A | 23 | 1 | 2.11* | ↓ in 7wk |
| Porphyrin Metabolism | 3.5A_7A | 31 | 5 | 2.01 | ↑ in 7wk |
|  | 3.5Ctrl_7Ctrl | 31 | 5 | 2.47 | ↑ in 7wk |
| Galactose Metabolism | 3.5A_3.5Ctrl | 27 | 1 | 2.35 | ↓ in Aden |
|  | 7A_7Ctrl | 27 | 5 | 2.07 | ↓ in Aden |
| Starch and sucrose metabolism | 7A_7Ctrl | 11 | 4 | 2.07 | ↓ in Aden |
| Steroid Hormone Biosynthesis | 3.5A_7A | 79 | 48 | 1.92 | ↑ in 7wk |
|  | 3.5Ctrl_7Ctrl | 79 | 48 | 2.39 | ↑ in 7wk |
| Taurine and Hypotaurine Metabolism | 3.5A_7A | 8 | 4 | 1.95 | ↑ in 7wk |
|  | 3.5Ctrl_7Ctrl | 8 | 4 | 2.58* | ↑ in 7wk |
| Ubiquinone and other terpenoid-quinone biosynthesis | 3.5A_7A | 18 | 5 | 1.97 | ↑ in 7wk |
|  | 3.5Ctrl_7Ctrl | 18 | 6 | 2.42 | ↑ in 7wk |
| Histidine Metabolism | 3.5A_3.5Ctrl | 16 | 1 | 2.19 | ↓ in Aden |
| Phenylalanine Metabolism | 3.5A_3.5Ctrl | 10 | 1 | 1.91 | ↓ in Aden |
| Tyrosine Metabolism | 3.5A_3.5Ctrl | 42 | 2 | 1.91 | ↓ in Aden |
|  | 7A_7Ctrl | 42 | 20 | 2.32 | ↑ in Aden |
| Cysteine and Methionine Metabolism | 3.5A_7A | 33 | 7 | 1.96 | ↓ in 7wk |
| Valine, Leucine, and Isoleucine Biosynthesis | M3.5A_M7A | 8 | 6 | 1.94* | ↓ in 7wk |
| Valine, Leucine, and Isoleucine Degradation | M3.5A_M7A | 40 | 9 | 1.94* | ↑ in 7wk |
| **(B) Pathways of interest from Volcano plots** |  |  |  |  |  |
| Amino sugar and nucleotide sugar metabolism | 3.5Ctrl_7Ctrl | 42 | 1 |  | ↑ in 7wk |
| Pentose Phosphate Pathway | 3.5A_7A | 23 | 1 |  | ↓ in 7wk |
| Porphyrin Metabolism | 3.5A_7A | 31 | 2 |  | ↑ in 7wk |
|  | 3.5Ctrl_7Ctrl | 31 | 2 |  | ↑ in 7wk |
| Taurine and Hypotaurine Metabolism | 3.5A_7A | 8 | 1 |  | ↑ in 7wk |
|  | 3.5Ctrl_7Ctrl | 8 | 1 |  | ↑ in 7wk |
| Ubiquinone and other terpenoid-quinone biosynthesis | 3.5A_7A | 18 | 2 |  | ↑ in 7wk |
|  | 3.5Ctrl_7Ctrl | 18 | 1 |  | ↑ in 7wk |
| Fatty Acid Degradation | 3.5A_7A | 39 | 1 |  | ↓ in 7wk |
| Biosynthesis of unsaturated fatty acids | 3.5Ctrl_7Ctrl | 36 | 1 |  | ↓ in 7wk |
| Steroid Biosynthesis | 3.5A_7A | 41 | 6 |  | ↑ in 7wk |
| Alanine, Aspartate, and Glutamate Metabolism | F3.5A_F7A | 28 | 1 |  | ↑ in 7wk |
| Arginine and Proline Metabolism | F3.5A_F7A | 36 | 1 |  | ↑ in 7wk |
| Glycine, Serine, and Threonine Metabolism | F3.5A_F7A | 34 | 1 |  | ↑ in 7wk |
| Cysteine and Methionine Metabolism | 3.5A_7A | 33 | 2 |  | ↓ in 7wk |
| Lysine Degradation | F3.5A_F7A | 30 | 3 |  | ↑ in 7wk |
| Valine, Leucine, and Isoleucine Biosynthesis | F3.5A_F7A | 8 | 3 |  | ↑ in 7wk |
| Valine, Leucine, and Isoleucine Degradation | F3.5A_F7A | 40 | 3 |  | ↑ in 7wk |

The VIP scores plot for the 3.5-vs 7-week adenine mice revealed that the top 100 metabolites had VIP scores between 1.87 and 2.27 (**Figure 5G**). Pathway analysis identified that mice with more severe CKD experience upregulations in some energy-related pathways (taurine/hypotaurine metabolism, ubiquinone and other terpenoid-quinone biosynthesis, porphyrin metabolism) while also experiencing downregulations in fatty acid degradation, pantothenate and CoA biosynthesis, the pentose phosphate pathway, and cysteine and methionine metabolism (**Table 5A**).

The comparison of mice fed a control diet over 3.5 and 7 weeks showed several similarities to the adenine comparison over the same time interval. From Volcano plot analysis (**Figure 5D**), the 296 metabolites found to be significantly dysregulated (214 upregulated and 82 downregulated in 7-week mice) were associated with upregulated amino sugar and nucleotide sugar metabolism, porphyrin metabolism, taurine and hypotaurine metabolism, and ubiquinone and other terpenoid-quinone biosynthesis (**Table 5B**). They were also associated with downregulated biosynthesis of unsaturated fatty acids (**Table 5B**). The top 100 metabolites (score range of 2.06-2.58) from the VIP score plot for the control diet comparison (**Figure 5H**) pertained to pathways also identified in the 3.5-versus 7-week adenine comparison (**Table 5A**). However, only the 3.5-vs 7-week adenine comparison identified dysregulated pentose phosphate pathway, cysteine metabolism, and methionine metabolism.

The effects of CKD severity on bone metabolism were further analyzed by comparing 3.5-week adenine and control mice along with 7-week adenine and control mice. 40.8% and 32.7% of the variation was accounted for in the PCA and PLS-DA plots for both comparisons, respectively (**Figure 5A, B**). The 3.5-week comparison produced 2 significant features from Volcano plot analysis that were downregulated in adenine mice, while the 7-week comparison identified none (**Figure 5E, F**). With only 2 features, the threshold value of metabolites was not achieved to permit pathway analysis. The top 100 metabolites for the 3.5-week comparison had VIP scores within the range of 1.87 and 3.91, while the range for the 7-week comparison was 1.96-2.96 (**Figure 5I, J**). Both comparisons experienced similarly downregulated pathways in adenine mice, most notably including galactose metabolism (**Table 5A**). Amino acid pathways were also impacted, including phenylalanine, tyrosine, and histidine metabolism. These amino acid pathways were downregulated in adenine vs control mice for the 3.5-week comparison, with tyrosine metabolism being upregulated in adenine vs control mice for the 7-week comparison.

## 4.0 Discussion

The purpose of this study was to determine the impact of 0.2% w/w adenine-induced kidney disease on cortical bone tissue metabolism and whether these impacts depend on disease severity (i.e., 3.5 vs 7 weeks of adenine) or male/female sex for C57BL/6J mice. We hypothesized that CKD would dysregulate bone tissue metabolism in a manner that depends on disease severity and sex, since these factors impact the response of C57BL/6 mice to an adenine diet as well as the extent of altered bone mass and quality.^28,35–38^ As expected, the adenine diet induced kidney disease in the study mice. At 3.5 and 7 weeks of diet, the mice had developed moderate and severe renal dysfunction, respectively (**Figure 2**). The males had more severe disease than females, as evidenced by ANOVA analyses of BUN and plasma creatinine. Our results are consistent with work by other groups found the adenine diet to induce a more severe disease in males than females.^38^ An untargeted metabolomics study was then conducted for marrow-flushed tibiae from these mice and comparisons made across factors of diet (adenine vs control), disease severity (3.5 weeks vs 7 weeks), and sex (female vs male).

We found that mice with adenine-induced CKD showed indications of altered cortical bone metabolism when compared to control mice. PCA and PLS-DA accounted for substantial variance and the top 100 VIP scores revealed several pathways that were altered in CKD. Adenine mice had downregulated essential (threonine metabolism, lysine degradation, and valine, leucine, and isoleucine biosynthesis and degradation) and nonessential (arginine, proline, glycine, and serine metabolism) amino acid pathways, fatty acid pathways, and pantothenate and CoA biosynthesis. Notably, metabolites associated with pantothenate and CoA biosynthesis, along with essential and nonessential amino acid pathways, all had VIP scores of 3.98, indicating the potential for them to serve as biomarkers of CKD.^43^ Amino acid metabolism is important for generating the components necessary for protein synthesis and bone anabolism managed by osteoblasts. ^44^ A downregulation in these pathways may suggest alterations to bone turnover, which is common in adenine-induced CKD.^28,35,37^ Furthermore, glutamate has been shown to impact bone cell (osteoblast and osteoclast) differentiation. ^45–49^ Fatty acid oxidation/degradation produces acetyl-CoA and is an important energy source for skeletal muscle, kidneys, and osteoblasts during bone building. ^50,51^ A downregulation in pantothenate and CoA biosynthesis could impact TCA cycle function and the synthesis of fatty acids, which could indicate alterations in energy use by bone cells. Together, these results suggest altered energy metabolism in cortical bone tissue resulting from CKD.

The impact of adenine-induced CKD on the cortical bone metabolome depended on diet length and thus the severity of disease. Compared to mice fed a 3.5-week adenine diet, the mice fed a 7-week adenine diet were found to have downregulated pentose phosphate pathway, pantothenate and CoA biosynthesis, fatty acid degradation, essential and nonessential amino acid pathways, and other energy-related pathways. These results were generally the same across volcano and top 100 VIP scores plots (**Figure 5C, G**). Many of the pathways that differed between 3.5-and 7-week adenine diet groups also differed between 3.5-and 7-week control diet groups. However, only the adenine diet comparison showed that the pentose phosphate pathway and cysteine and methionine metabolism were dysregulated with diet length (**Figure 5D, H**). The pentose phosphate pathway is a process that creates precursors for nucleotide and amino acid biosynthesis as well as provides reducing molecules for anabolic processes.^52^ A dysregulation in this pathway could be indicative of alterations in energy usage by bone cells.

To further assess the impact of CKD severity on bone metabolism, we then compared the 3.5-week and 7-week adenine diet groups to their respective controls. The top 100 VIP scores plots identified that amino acid (i.e., phenylalanine, tyrosine, and histidine metabolism) and galactose metabolism differed in expression between adenine-treated mice and their controls at both diet time points (**Figure 5I, J**). 7-week adenine mice also experienced dysregulated starch and sucrose metabolism compared to their controls, a trend not observed in the 3.5-week comparison. Discrepancies in essential/nonessential amino acid pathways and sugar metabolism may suggest that bone cell function is altered, with cells potentially utilizing energy differently in CKD-affected mice. Moreover, the expression of starch and sucrose metabolism exclusively in the 7-week adenine vs control comparison may indicate that bone cell energy utilization changes with CKD progression.

The impact of adenine-induced CKD on cortical bone metabolism was sexually dimorphic. These results are in line with prior studies that reported that changes to bone properties in C57BL/6 mice (with CKD) are also dependent on sex (**Table 1**).^28^ The top 100 VIP scores revealed that both male and female adenine mice had downregulated pantothenate and CoA biosynthesis, essential and nonessential amino acid pathways, and purine metabolism compared with control diet mice (**Figure 4G, H**). Downregulated purine metabolism is counterintuitive for the adenine model but, importantly, represents a single timepoint of measurement and does not capture the disease progression. Several pathways were unique to each sex. In males, the adenine diet affected fatty acid pathways while in females, the adenine diet dysregulated sugar pathways, porphyrin metabolism, and histidine metabolism. Female adenine mice were also compared with male adenine mice (**Figure 4J**). Female adenine mice had upregulated pantothenate and CoA biosynthesis among other energy-relevant pathways (i.e., ubiquinone and terpenoid-quinone biosynthesis), compared to male adenine mice. Pantothenate and CoA biosynthesis, along with ubiquinone and terpenoid-quinone biosynthesis, operate in concert with the TCA cycle and the electron transport chain (ETC), with ubiquinone being important for electron transport in the ETC.^53^ Alterations in ubiquinone biosynthesis could indicate that energy is being derived in an altered manner.

Control diet mice also showed sexually dimorphic alterations in cortical bone metabolism. These findings are complementary to prior studies that found C57BL/6 mice to have sex differences in bone properties and metabolism. ^21,22,30–34^ From the top 100 VIP scores, control female mice showed upregulated pantothenate and CoA biosynthesis, galactose metabolism, and essential and nonessential amino acid pathways compared to control males (**Figure 4I**). Female mice also had downregulations in fatty acid degradation, steroid hormone biosynthesis, tyrosine metabolism, and ubiquinone and other terpenoid-quinone biosynthesis. Alterations in these pathways may be indicative of differences in bone turnover and bone cell energy utilization between sexes.^44–51^ Galactose, starch, and sucrose metabolism were only observed in the control sex comparison (upregulated in females), while fatty acid elongation was only observed in the adenine sex comparison (downregulated in females).

Our work contributes to a small but growing body of work reporting sex differences in the metabolomes of cortical bone tissue. Other studies from our group have investigated sex differences in cortical bone tissue, albeit from humeri and different ages (i.e., 5 months) and strains (i.e., C57BL/6N and C57BL/6JN) of mice.^20–22^ The present study and the prior studies all found an amino acid pathway (tyrosine) to be upregulated in control males versus females. However, in this study but not the others, the remaining amino acid pathways were upregulated in females compared to males. Discrepancies in lipid metabolism were also observed between studies. Each of these studies have identified alterations in lipid metabolism between sexes, but the direction is not always consistent. Pathways found to differ between sexes in the prior studies, but not this one, included aspartate, glutamate, cysteine, and methionine metabolism along with glycolysis/gluconeogenesis and the TCA cycle. Pathways that were observed in this study and not the others included starch, sucrose, and galactose metabolism, pantothenate and CoA biosynthesis, glycine, serine, and threonine metabolism, along with valine, leucine, and isoleucine biosynthesis and degradation. These pathways were all upregulated in control females versus males.

These differences in pathway expression could be at least partially attributable to the mice being raised at Georgetown University as opposed to Montana State University. Factors associated with how they were raised (temperature, diet, elevation, etc.) and their breeding/sourcing could have also contributed to the observed differences.

This study has several important limitations. Additional bone tissues were not available for bone quality analyses; however, bone quality changes in the adenine model of disease has been well-characterized in C57BL/6 mice for this range of treatment times (**Table 1**).^28,35–38^ While our procedure for flushing marrow from long bones is successful in removing marrow tissues, it is possible that small capillaries and residual metabolites persist within the bone tissue, which implies that not all metabolites are produced by osteocytes.

In summary, this study investigated the impacts of adenine-induced CKD, disease severity, and sex on the cortical bone metabolome. These data suggest that altered cellular bioenergetics may have a role in the decreased bone quality in CKD and highlight several potential biomarkers worthy of additional investigation. Further, the data provide evidence that sex differences in bone health extend to the level of bone tissue metabolism. Collectively, these results motivate further inquiry into the role of bone tissue metabolism in the progressive loss of bone fracture resistance in the context of CKD.

## Acknowledgements

The authors gratefully acknowledge support from Montana State University’s Department of Mechanical and Industrial Engineering and the Norm Asbjornson College of Engineering. Research reported in this publication was supported by the National Institute of General Medical Sciences of the National Institutes of Health under Award Number P20GM103474 and the National Science Foundation award 2340823. The content is solely the responsibility of the authors and does not necessarily represent the official views of the sponsors.

