## Supplementary figures and tables for "Adenine-induced kidney disease alters the cortical bone metabolome of C57BL/6J mice in a manner that depends on sex"

**Supplementary Information**

***ECCO Analysis***

ECCO was used to perform clustering analyses for all samples (i.e., metabolites of all mice) in the study. From the original 3546 metabolites identified from LC-MS, ECCO identified 3167 metabolites that clustered together 100% of the time based on an ensemble that employed 13 clustering solutions. Single, complete, and average linkage functions were applied to Euclidean, square root Euclidean, standardized Euclidean, and Chebyshev distance functions. The ward linkage function was also applied to the Euclidean distance function, bringing the total number of clustering solutions to 13. Two clusters were identified during ECCO's clustering optimization step (430 clusters and 2737 metabolites in each cluster). These clusters did not strongly align with any of the three factors compared in this study: CKD, sex, and severity (**Figure S.1**).

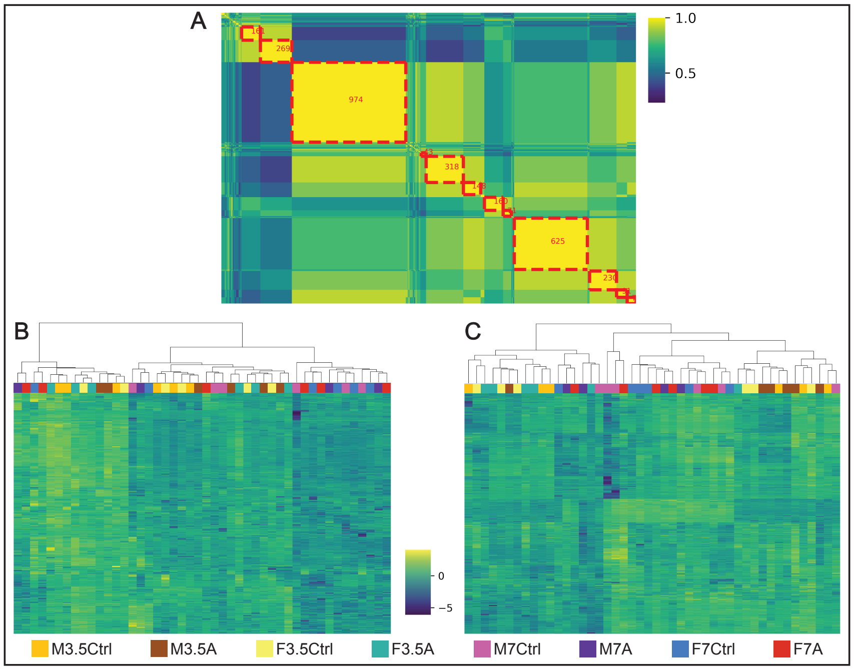


**Figure S.1.** Metabolite ensemble outputs from ECCO. (A) Metabolite ensemble output, with coloration ranging from purple to yellow being indicative of clustering percentage across multiple clustering algorithms. Regions of yellow that are boxed in red are metabolites that clustered together in

100% of clustering solutions, and percentage decreases as the region's color gets closer to purple. (B) The heatmap of the expression of cluster 1 metabolites within all study samples. Cluster 1 is the region of green and yellow in the top left corner of the metabolite ensemble output. The metabolites that were found to be clustering together within cluster 1 in the metabolite ensemble output were identified in all of the samples within the study. Hierarchical clustering was used to group samples based on similar expression of cluster 1 metabolites, with the groups that samples were a part of being identified on the top row of the plot. Minimal group-specific clustering was determined from the heatmap of cluster 1 metabolites from the metabolite ensemble result produced by ECCO. (C) The heatmap of the expression of cluster 2 metabolites within all study samples. Cluster 2 is the much larger region of green and yellow in the metabolite ensemble output, with it ranging from the box of 974 metabolites adjacent to cluster 1 to the bottom right corner of the plot. The heatmap associated with cluster 2 was produced and analyzed in a very similar manner to the heatmap of cluster 1. Similarly to cluster 1, no group-specific clustering was observed based on the expression of cluster 2 metabolites.

| Source | DF | Adj SS | Adj MS | F-Value | P-Value |
|---|---|---|---|---|---|
| Sex | 1 | 513.4 | 513.4 | 12.02 | 0.001 |
| Duration | 1 | 744.7 | 744.7 | 17.43 | <0.001 |
| Diet | 1 | 11557.6 | 11557.6 | 270.54 | <0.001 |
| Sex*Duration | 1 | 46.1 | 46.1 | 1.08 | 0.306 |
| Sex*Diet | 1 | 152.4 | 152.4 | 3.57 | 0.067 |
| Duration*Diet | 1 | 1144.4 | 1144.4 | 26.79 | <0.001 |
| Sex*Duration*Diet | 1 | 63.1 | 63.1 | 1.48 | 0.232 |
| Error | 38 | 1623.4 | 42.7 | | |
| Total | 45 | 14796.7 | | | |

**Table S.1.** Results from 3-way ANOVA of BUN levels.

| Source | DF | Adj SS | Adj MS | F-Value | P-Value |
| --- | --- | --- | --- | --- | --- |
| Sex | 1 | 0.2478 | 0.2478 | 10.56 | 0.002 |
| Duration | 1 | 0.1106 | 0.1106 | 4.71 | 0.036 |
| Diet | 1 | 1.42167 | 1.42167 | 60.59 | <0.001 |
| Sex*Duration | 1 | 0.00036 | 0.00036 | 0.02 | 0.902 |
| Sex*Diet | 1 | 0.01271 | 0.01271 | 0.54 | 0.466 |
| Duration*Diet | 1 | 0.0828 | 0.0828 | 3.53 | 0.068 |
| Sex*Duration*Diet | 1 | 0.00338 | 0.00338 | 0.14 | 0.707 |
| Error | 38 | 0.89159 | 0.02346 | | |
| Total | 45 | 2.64933 | | | |

**Table S.2.** Results from 3-way ANOVA of plasma creatinine levels.

| Pathway | Comparison | Pathway Total | Pathway Hits | VIP Score | Trends |
|---|---|---|---|---|---|
| Fatty Acid Biosynthesis | F3.5A_F3.5Ctrl | 47 | 3 | 2.09 | ↓ in Aden |
| | M3.5A_M3.5Ctrl | 47 | 2 | 2.36 | ↑ in Aden |
| Fatty Acid Degradation | F3.5A_F3.5Ctrl | 39 | 8 | 2.09 | ↓ in Aden |
| | F7A_F7Ctrl | 39 | 4 | 2.14 | ↓ in Aden |
| Fatty Acid Elongation | F3.5A_F3.5Ctrl | 39 | 6 | 2.09 | ↓ in Aden |
| Steroid Hormone Biosynthesis | F3.5A_F3.5Ctrl | 79 | 53 | 2.21 | ↓ in Aden |
| | M3.5A_M3.5Ctrl | 79 | 52 | 2.16 | ↑ in Aden |
| | M7A_M7Ctrl | 79 | 47 | 2.21 | ↑ in Aden |
| Galactose Metabolism | F3.5A_F3.5Ctrl | 27 | 5 | 1.85 | ↓ in Aden |
| | F7A_F7Ctrl | 27 | 5 | 2.64* | ↑ in Aden |
| | M3.5A_M3.5Ctrl | 27 | 5 | 1.92 | ↑ in Aden |
| Starch and Sucrose Metabolism | F3.5A_F3.5Ctrl | 15 | 4 | 1.85 | ↓ in Aden |
| | F7A_F7Ctrl | 15 | 4 | 2.64* | ↑ in Aden |
| | M3.5A_M3.5Ctrl | 15 | 4 | 1.92 | ↑ in Aden |
| Pantothenate and CoA Biosynthesis | F3.5A_F3.5Ctrl | 20 | 3 | 2.53* | ↓ in Aden |
| | M3.5A_M3.5Ctrl | 20 | 2 | 2.61 | ↓ in Aden |
| | M7A_M7Ctrl | 20 | 2 | 2.31* | ↓ in Aden |
| Tyrosine Metabolism | F3.5A_F3.5Ctrl | 42 | 20 | 2.12 | ↓ in Aden |
| | F7A_F7Ctrl | 42 | 20 | 2.36 | ↓ in Aden |
| | M3.5A_M3.5Ctrl | 42 | 18 | 2.57 | ↑ in Aden |
| | M7A_M7Ctrl | 42 | 17 | 1.89 | ↓ in Aden |
| Alanine, Aspartate, and Glutamate Metabolism | F3.5A_F3.5Ctrl | 28 | 2 | 1.85 | ↓ in Aden |
| | F7A_F7Ctrl | 28 | 2 | 2.03 | ↓ in Aden |
| | M3.5A_M3.5Ctrl | 28 | 2 | 1.97 | ↑ in Aden |
| Arginine and Proline Metabolism | F3.5A_F3.5Ctrl | 36 | 8 | 2.53* | ↓ in Aden |
| | M3.5A_M3.5Ctrl | 36 | 8 | 2.61 | ↓ in Aden |
| | M7A_M7Ctrl | 36 | 8 | 2.31* | ↓ in Aden |
| Glycine, Serine, and Threonine Metabolism | F3.5A_F3.5Ctrl | 34 | 3 | 2.53* | ↓ in Aden |
| | F7A_F7Ctrl | 34 | 3 | 2.03 | ↓ in Aden |
| | M3.5A_M3.5Ctrl | 34 | 3 | 2.61 | ↑ in Aden |
| | M7A_M7Ctrl | 34 | 3 | 2.31* | ↓ in Aden |
| Lysine Degradation | F3.5A_F3.5Ctrl | 30 | 7 | 2.53* | ↓ in Aden |
| | M3.5A_M3.5Ctrl | 30 | 6 | 2.61 | ↓ in Aden |
| | M7A_M7Ctrl | 30 | 6 | 2.31* | ↓ in Aden |
| Valine, Leucine, and Isoleucine Biosynthesis | F3.5A_F3.5Ctrl | 8 | 6 | 2.53* | ↓ in Aden |
| | F7A_F7Ctrl | 8 | 6 | 2.03 | ↓ in Aden |
| | M3.5A_M3.5Ctrl | 8 | 6 | 2.61 | ↓ in Aden |
| | M7A_M7Ctrl | 8 | 6 | 2.31* | ↓ in Aden |
| Valine, Leucine, and Isoleucine Degradation | F3.5A_F3.5Ctrl | 40 | 10 | 2.53* | ↓ in Aden |
| | F7A_F7Ctrl | 40 | 9 | 2.03 | ↓ in Aden |
| | M3.5A_M3.5Ctrl | 40 | 9 | 2.61 | ↓ in Aden |
| | M7A_M7Ctrl | 40 | 9 | 2.31* | ↓ in Aden |

**Table S.3.** Metabolic pathways identified in comparisons between mice with and without adenine-induced CKD. VIP scores labeled with an asterisk indicate the pathways that were most significantly dysregulated in their associated comparisons.

| Pathway | Comparison | Pathway Total | Pathway Hits | VIP Score | Trends |
|---|---|---|---|---|---|
| Fatty Acid Degradation | F3.5Ctrl_M3.5Ctrl | 39 | 7 | 2.22 | ↓ in Females |
| | F7A_M7A | 39 | 5 | 1.95 | ↑ in Females |
| Fatty Acid Elongation | F3.5Ctrl_M3.5Ctrl | 39 | 6 | 2.06 | ↑ in Females |
| | F7A_M7A | 39 | 4 | 1.95 | ↑ in Females |
| Fatty Acid Biosynthesis | F3.5Ctrl_M3.5Ctrl | 47 | 3 | 2.06 | ↑ in Females |
| Biosynthesis of Unsaturated Fatty Acids | F3.5Ctrl_M3.5Ctrl | 36 | 9 | 2.09 | ↑ in Females |
| Drug Metabolism – cytochrome P450 | F3.5Ctrl_M3.5Ctrl | 27 | 7 | 2.63 | ↓ in Females |
| | F7Ctrl_M7Ctrl | 27 | 7 | 1.9 | ↓ in Females |
| Metabolism of Xenobiotics by Cytochrome P450 | F3.5Ctrl_M3.5Ctrl | 64 | 2 | 2.88* | ↑ in Females |
| | F7Ctrl_M7Ctrl | 64 | 2 | 2.36 | ↓ in Females |
| Galactose Metabolism | F3.5Ctrl_M3.5Ctrl | 27 | 5 | 2.51 | ↑ in Females |
| | F7Ctrl_M7Ctrl | 27 | 5 | 2.41 | ↑ in Females |
| Starch and Sucrose Metabolism | F3.5Ctrl_M3.5Ctrl | 15 | 4 | 2.51 | ↑ in Females |
| | F7Ctrl_M7Ctrl | 15 | 4 | 2.41 | ↑ in Females |
| Pantothenate and CoA Biosynthesis | F7Ctrl_M7Ctrl | 20 | 2 | 2.33 | ↑ in Females |
| | F7A_M7A | 20 | 2 | 2.86* | ↑ in Females |
| Ubiquinone and other terpenoid-quinone biosynthesis | F7A_M7A | 18 | 5 | 2.06 | ↑ in Females |
| Steroid Hormone Biosynthesis | F3.5Ctrl_M3.5Ctrl | 79 | 54 | 2.01 | ↑ in Females |
| | F7Ctrl_M7Ctrl | 79 | 46 | 2.45* | ↑ in Females |
| | F3.5A_M3.5A | 79 | 53 | 1.93 | ↑ in Females |
| | F7A_M7A | 79 | 50 | 2.5 | ↑ in Females |
| Tyrosine Metabolism | F3.5Ctrl_M3.5Ctrl | 42 | 19 | 2.13 | ↑ in Females |
| | F7A_M7A | 42 | 20 | 2.19 | ↑ in Females |
| Tryptophan Metabolism | F3.5Ctrl_M3.5Ctrl | 41 | 2 | 2.06 | ↑ in Females |
| Arginine and Proline Metabolism | F7Ctrl_M7Ctrl | 36 | 8 | 2.24 | ↑ in Females |
| | F7A_M7A | 36 | 8 | 2.86* | ↑ in Females |
| Glycine, Serine, and Threonine Metabolism | F7Ctrl_M7Ctrl | 34 | 3 | 2.33 | ↑ in Females |
| | F7A_M7A | 34 | 3 | 2.86* | ↑ in Females |
| Lysine Degradation | F3.5Ctrl_M3.5Ctrl | 30 | 7 | 2.06 | ↑ in Females |
| | F7Ctrl_M7Ctrl | 30 | 6 | 2.24 | ↑ in Females |
| | F7A_M7A | 30 | 6 | 2.86* | ↑ in Females |
| Valine, Leucine, and Isoleucine Biosynthesis | F7Ctrl_M7Ctrl | 8 | 6 | 2.33 | ↑ in Females |
| Valine, Leucine, and Isoleucine Degradation | F3.5Ctrl_M3.5Ctrl | 40 | 10 | 2.06 | ↑ in Females |
| | F7Ctrl_M7Ctrl | 40 | 9 | 2.33 | ↑ in Females |

**Table S.4.** Metabolic pathways identified in comparisons between male and female mice. VIP scores labeled with an asterisk indicate the pathways that were most significantly dysregulated in specific comparisons.

| Pathway | Comparison | Pathway Total | Pathway Hits | VIP Score | Trends |
|---|---|---|---|---|---|
| **(A) Pathways of interest from top 100 VIP scores** | | | | | |
| Amino sugar and nucleotide sugar metabolism | M3.5Ctrl_M7Ctrl | 42 | 2 | 2.22 | ↑ in 7wk |
| Pentose Phosphate Pathway | F3.5A_F7A | 23 | 1 | 1.75 | ↓ in 7wk |
| | M3.5A_M7A | 23 | 1 | 1.79 | ↓ in 7wk |
| | F3.5Ctrl_F7Ctrl | 23 | 1 | 2.19 | ↑ in 7wk |
| Porphyrin Metabolism | F3.5A_F7A | 31 | 5 | 1.77 | ↑ in 7wk |
| | F3.5Ctrl_F7Ctrl | 31 | 5 | 2.28* | ↓ in 7wk |
| | M3.5Ctrl_M7Ctrl | 31 | 5 | 2.19 | ↑ in 7wk |
| Steroid Hormone Biosynthesis | F3.5A_F7A | 79 | 51 | 1.76 | ↑ in 7wk |
| | M3.5A_M7A | 79 | 51 | 1.84 | ↑ in 7wk |
| | F3.5Ctrl_F7Ctrl | 79 | 47 | 2.2 | ↑ in 7wk |
| Taurine and Hypotaurine Metabolism | F3.5A_F7A | 8 | 4 | 1.86* | ↑ in 7wk |
| | M3.5A_M7A | 8 | 4 | 1.79 | ↑ in 7wk |
| | F3.5Ctrl_F7Ctrl | 8 | 4 | 2.24 | ↑ in 7wk |
| | M3.5Ctrl_M7Ctrl | 8 | 4 | 2.43* | ↑ in 7wk |
| Ubiquinone and other terpenoid-quinone biosynthesis | F3.5A_F7A | 18 | 5 | 1.76 | ↑ in 7wk |
| | F3.5Ctrl_F7Ctrl | 18 | 5 | 2.19 | ↑ in 7wk |
| Cysteine and Methionine Metabolism | M3.5A_M7A | 33 | 7 | 1.77 | ↓ in 7wk |
| Valine, Leucine, and Isoleucine Biosynthesis | M3.5A_M7A | 8 | 6 | 1.94* | ↓ in 7wk |
| | M3.5Ctrl_M7Ctrl | 8 | 6 | 2.16 | ↑ in 7wk |
| Valine, Leucine, and Isoleucine | M3.5A_M7A | 40 | 9 | 1.94* | ↑ in 7wk |
| | M3.5Ctrl_M7Ctrl | 40 | 9 | 2.16 | ↑ in 7wk |
| **(B) Pathways of interest from Volcano plots** | | | | | |
| Pentose Phosphate Pathway | F3.5A_F7A | 23 | 1 | | ↓ in 7wk |
| Porphyrin Metabolism | F3.5A_F7A | 31 | 2 | | ↑ in 7wk |
| Taurine and Hypotaurine Metabolism | F3.5A_F7A | 8 | 1 | | ↑ in 7wk |
| Ubiquinone and other terpenoid-quinone biosynthesis | F3.5A_F7A | 18 | 1 | | ↑ in 7wk |
| Fatty Acid Degradation | F3.5A_F7A | 39 | 1 | | ↓ in 7wk |
| Steroid Biosynthesis | F3.5A_F7A | 41 | 7 | | ↑ in 7wk |
| Alanine, Aspartate, and Glutamate Metabolism | F3.5A_F7A | 28 | 1 | | ↑ in 7wk |
| Arginine and Proline Metabolism | F3.5A_F7A | 36 | 1 | | ↑ in 7wk |
| Glycine, Serine, and Threonine Metabolism | F3.5A_F7A | 34 | 1 | | ↑ in 7wk |
| Cysteine and Methionine Metabolism | F3.5A_F7A | 33 | 2 | | ↓ in 7wk |
| Lysine Degradation | F3.5A_F7A | 30 | 3 | | ↑ in 7wk |
| Valine, Leucine, and Isoleucine Biosynthesis | F3.5A_F7A | 8 | 3 | | ↑ in 7wk |
| Valine, Leucine, and Isoleucine Degradation | F3.5A_F7A | 40 | 3 | | ↑ in 7wk |

**Table S.5.** Metabolic pathways identified in comparisons of 3.5 or 7 weeks of adenine or control diets. (A) Pathways identified from top 100 VIP scores plots for each of the relevant comparisons. (B) Pathways identified from Volcano plots for each of the relevant comparisons. VIP scores with an asterisk indicate the pathways that were most significantly dysregulated in specific comparisons.
